# Using Manganese Enhanced Magnetic Resonance Imaging (MEMRI) to Assess Calcium Dependent Activity in Vestibular Pathways Following Linear Acceleration

**DOI:** 10.1101/2023.11.29.569068

**Authors:** Danial Naqvi, Rod D. Braun, Antonela Muca, Mirabela Hali, Avril Genene Holt

## Abstract

Reliable methods for repetitive and longitudinal assessment of central vestibular pathway function *in vivo* are rather limited. Manganese-enhanced magnetic resonance imaging (MEMRI) has been used in various sensory systems to evaluate neuronal activity in central pathways, but MEMRI assessment of central vestibular pathways has been minimal. The present study addressed this gap in knowledge by assessing whether Mn^2+^ can be taken up in an activity-dependent manner through voltage-gated calcium channels in the vestibular nuclear complex (VNC) and the vestibulocerebellum (VeCb) of rats with and without mild linear acceleration stimulation. *R*1 maps were collected prior to, one day after, and two weeks after Mn^2+^ administration in stimulated and non-stimulated rats. Analysis of MRI *R*1 values showed that one day after Mn^2+^ administration the VNC and VeCb had significantly greater *R*1 values that returned to baseline levels after two weeks. Non-stimulated rats had greater *R*1 values than stimulated rats. Mid rostro-caudal sections of the VNC had greater *R*1 values than rostral and caudal VNC sections. *R*1 values also indicated that Mn^2+^ was differentially taken up across subdivisions of the VNC and VeCb. These results correlate well with expected patterns of neuronal activity after linear acceleration. MEMRI is a sensitive tool that may be used to evaluate activity patterns in central vestibular nuclei, proving useful for studying underlying mechanisms of central vestibular dysfunction.

## 1. Introduction

Manganese (Mn^2+^)-enhanced magnetic resonance imaging (MEMRI) is often used as an *in vivo* measure of central activity (Lin and Koretsky, 1997; Pautler et al., 1998; Silva et al., 2004; Pautler, 2006). Assessment of c-Fos production has long been the “gold standard” for measurement of neuronal activity (Morgan et al., 1987; Sagar et al., 1988), but this approach does not traditionally allow longitudinal assessment. In contrast, MEMRI allows repetitive *in vivo* measurement of central activity. As a paramagnetic agent, Mn^2+^ shortens T1 relaxation times, providing contrast enhancement of T1-weighted images and T1 Maps (Mendonça-Dias et al., 1983; Geraldes et al., 1986; Cory et al., 1987; Fornasiero et al., 1987). In addition, Mn^2+^ acts as a calcium surrogate in neurons (Narita et al., 1990; Drapeau and Nachshen, 1984), entering through voltage- and ligand-gated calcium channels (Kita et al., 1981; Burnett et al., 1984; Cross et al., 2007; Berkowitz et al., 2007; Gadjanski et al., 2009; Finley, 1998). Thus Mn^2+^ acts as an activity-dependent contrast agent. Changes in neuronal activity result in more or less Mn^2+^ accumulation in the cell, relative to that measured under control conditions (Itoh et al., 2008). Slow efflux from cells allows for assessment of Mn^2+^ uptake from several hours to days after the activating event (Aoki et al., 2004). The use of MEMRI in auditory pathways has been demonstrated in animal models of noise induced hearing loss (NIHL) and tinnitus (Yu et al., 2005; Yu et al., 2008; Brozoski et al., 2007; Holt et al., 2010; Brozoski et al., 2013; Cacace et al., 2014; Muca et al., 2018). Although MEMRI has been used to assess neural activity in various sensory systems, such as olfactory, visual, and auditory systems (Tang et al., 2016; Pautler and Koretsky, 2002; Tjalve et al., 1995; Tjalve et al., 1996; Bissig and Berkowitz, 2014), studies of the vestibular and balance pathways using MEMRI have been limited (Rodriguez et al., 2016). Providing evidence that Mn^2+^ can be taken up by central vestibular-related neurons would prove useful for future longitudinal studies.

All afferents innervating peripheral vestibular end-organs send projections to either the vestibular nuclear complex (VNC) or the vestibulocerebellum (VeCb; Maklad et al., 2010; Eatock, 2008). Within the sensory epithelium of each vestibular end-organ is a central region containing type I sensory hair cells. In the otolith organs (the saccule and utricle) this central region, the striola, is innervated by calyx-only ganglion cells (irregular afferent fibers) (Desmadryl and Dechesne, 1992; Eatock et al., 2008; Goldberg et al., 1990; Curthoys et al., 2017; Eatock 2018). These irregular fibers detect rapid linear head motion and therefore are responsive to a jerk stimulus. Within the extra-striolar (peripheral) zone are type II hair cells innervated by bouton-only terminals (regular afferent fibers) that are responsible for detecting slow head motion. The third type of afferents are dimorphic, which innervate type I and type II hair cells found within the striolar and extrastriolar regions (Fernández et al., 1990). The vestibulocochlear nerve comprises projections from each of these afferents, which form the vestibular portion of CN VIII, and synapse with neurons within the VNC and VeCb.

Otolith irregular afferents have enriched projections to specific VNC and VeCb nuclei. The VNC consists of four major nuclei: lateral (LVe), medial (MVe), superior (SuVe), and inferior vestibular nuclei (spinal vestibular nucleus in rats - SpVe). These VNC nuclei are found along the floor of the fourth ventricle, spanning from the rostral medulla to the caudal pons, and receive ipsilateral projections from CN VIII (Barmack, 2003). The VeCb consists of the flocculus, nodulus (lobule X), ventral-paraflocculus, and uvula (lobule XI). The flocculus and paraflocculus are located in the cerebellar pontine angle in humans which is analogous to the posterior border of the middle cerebellar peduncle in the rat (Mennink et al., 2020). The uvula and nodulus (lobules IX and X, respectively) are found in the most posterior portion of the cerebellar vermis. They receive ipsilateral projections from CN VIII (Voogd et al., 1996). While afferents from the periphery carry signals from specific vestibular end organs to neurons throughout the VNC, there are some distinctions. Canal afferents terminate primarily in the SuVe and MVe while otolith afferents primarily innervate the LVe and SpVe nuclei (Newlands and Perachio, 2003; Büttner-Ennerver, 1992). The SuVe and MVe nuclei play important roles in control of eye movement (Vidal and Sans, 2004) while the LVe and SpVe nuclei have descending projections to extensor muscles and affect autonomic function (Holstein et al., 2011; Barmack, 2003). We expect each of these VNC subdivisions to take up Mn^2+^ due to their spontaneous neuronal activity, but we expect otolith stimulation to result in differential uptake in the LVe and SpVe nuclei. Therefore, we administer Mn^2+^ and map the resulting uptake by VNC nuclei in non-stimulated and stimulated animals. To begin to unravel the questions of causality we test the effect of mild peripheral stimulation on activation of central nuclei using Mn^2+^ uptake as an *in vivo* marker of neural activity. These studies lay the foundation for future *in vivo* longitudinal imaging studies in vestibular pathways.

## 2. Materials and Methods

### 2.1. Subjects and design

All procedures were approved by the Institutional Animal Care and Use Committee (IACUC) at Wayne State University. Male Sprague-Dawley rats (300 – 400 grams) were obtained from Charles River Laboratory. All rats were individually housed and maintained at 77°F with a 12 – hour light/dark cycle (lights on at 7 AM). Standard housing conditions with free access to normal rat chow and tap water were provided. Wayne State University (WSU) maintains AAALAC accredited animal facilities under the Division of Laboratory Animal Resources (DLAR). After arrival at the WSU DLAR facility, animals were allowed to acclimate for at least 24 hours before undergoing surgical procedures and the ten rats were divided into two groups: a sham group (n=5) and a stimulation group (n=5). All rats underwent surgery (described below), to place a nut or head bolt on the skull, which permitted attachment of the head to a shaker. Ten days after surgery, rats underwent MRI imaging to obtain baseline *R*1 maps of the VNC and VeCb of the brain. Two to three days later, rats in both groups were administered MnCl_2_, and while the rats in the sham group were attached to the shaker, they were not stimulated. However, rats in the stimulated group were attached to the shaker and were delivered the jerk stimulus paradigm (see below). After MnCl_2_ administration (16-24 hours), all rats were imaged (MEMRI). Rats from each group were imaged again two weeks after MnCl_2_ administration to assess Mn^2+^ clearance.

### 2.2. Surgical procedures

All surgical procedures were performed under aseptic conditions. Animals were anesthetized with ketamine (75 mg/kg) and xylazine (8 mg/kg) administered intraperitoneally (i.p.). After stabilization and leveling of the head using a stereotaxic frame, bregma and lambda were exposed, and two anchor screws were placed into the skull. A 6-32 HEX M/S ceramic nut (Small Parts Inc., model #B000FN0BZG, UNPSC code 31161700) or a custom-made titanium head bolt was centered on bregma and affixed to the skull using C&B Metabond cement (Parkell, Edgewood, NY). The ceramic nut or head bolt was further secured to the skull using dental acrylic (Lang Dental Manufacturing Co., Inc. Wheeling, IL). The animals were given 7-10 days to recover prior to additional experiments (Fig. 1A).

**Figure 1.**
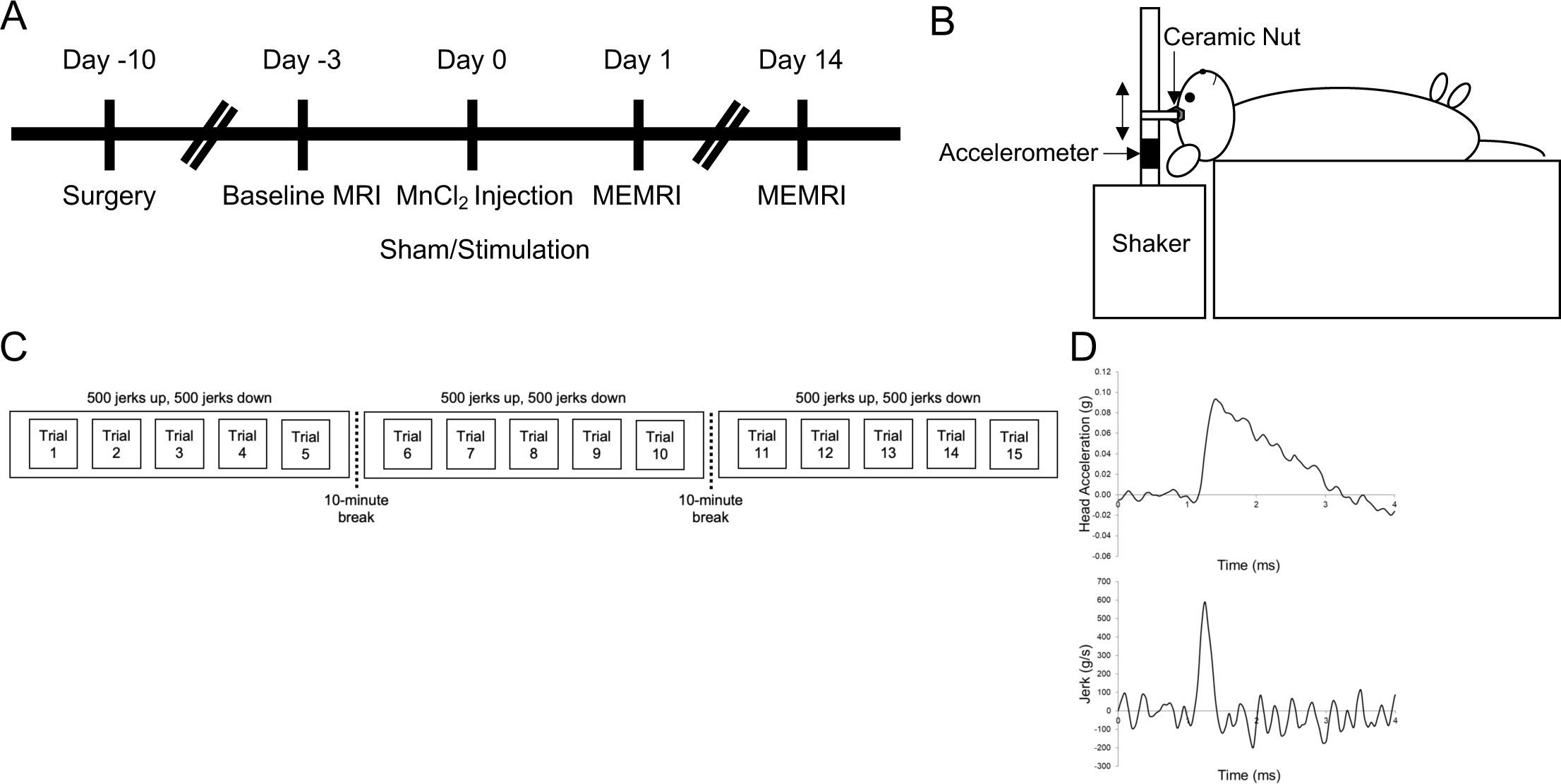
Experimental design and jerk paradigm. Imaging was conducted one day prior to (n=4), one day after (n=10), and two weeks after (n=4) Mn^2+^ administration (A). Prior to imaging, a ceramic nut was surgically affixed to the skull, allowing the head to be attached to the mechanical shaker (B). Stimulated rats were exposed to a total of 15 trials, split into three blocks of five trials each (C). Each trial consisted of 100 jerks up and 100 jerks down for a total of 3,000 jerks across all trials. Each block was followed by a 10-minute quiescent interval (break). Rats were stimulated in the naso-occipital plane at a jerk intensity of ∼ 600 g/s (D). Sham rats were attached to the shaker for the same duration but were not stimulated. Double-headed arrow in panel B depicts direction of jerk stimulation.

### 2.3. Mn^2+^ administration and stimulation of otolith organs

On the day of testing, animals were anesthetized with an intramuscular injection of ketamine (75 mg/kg) and xylazine (8 mg/kg) and placed in a supine position. Needle electrodes were positioned on the vertex of the skull (active), right ear (reference), and hip (ground). The electrode signals were filtered, digitized, and amplified using Spike2 software and CED Power 1401 data acquisition system at 200 kHz. The head was attached to a mechanical shaker arm (via bolt or ceramic nut) fitted with an accelerometer (Fig. 1B). Each animal was administered manganese (II) chloride tetrahydrate (MnCl_2;_ 66 mg/kg ip, Sigma Aldrich Co., St. Louis, MD) after attachment to the shaker arm. The sham group was subjected to the same conditions as the stimulated group but without delivery of the jerk stimulation. Each sham animal remained on the shaker platform for a period of 30 minutes. Stimulated rats were exposed to a total of 15 trials, split across three blocks comprising five trials each (Fig. 1C). A series of 200 jerks (100 up and 100 down) was designated as a trial. A 10-minute rest period was introduced after each block. The stimulation generated by the mechanical shaker had an amplitude of ∼ 600 g/s (Fig. 1D).

### 2.4. Manganese-enhanced magnetic resonance imaging (MEMRI)

During each imaging session, rats were anesthetized by inhalation of isoflurane (3 - 4% for induction and 1 - 1.5% for maintenance) mixed with room air and were placed on a heated re-circulating waterbed. The isoflurane was delivered through a nose cone and the head was stabilized by a bite bar. The regions of interest were imaged using a 7T MRI scanner (Bruker ClinScan) with a Siemens console (Siemens Corporation, Washington DC, USA). Prior to proceeding with the image acquisition, T_1_ weighted MRI localizer images were obtained to allow proper orientation of the brain (Muca et al., 2018).

### 2.5. T1 mapping

The Paxinos and Watson rat atlas (2007) was used to identify the location of vestibular nuclei in T1 maps. Six coronal, 400 µm, T1 map sections, were collected serially throughout the rostro-caudal extent of the VNC. In addition, one T1 map section was collected that encompassed the mid rostro-caudal inferior colliculus (encompassing the PnO - oral part of the pontine reticular nucleus and the Ent - entorhinal cortex). The most rostral VNC section was imaged by locating the most prominent occurrence of CN VIII and the cochlea in the same dorso-ventral plane and collecting the 400 µm section immediately caudal. Then, five additional serial sections were collected moving caudally, in 400 µm increments. The inferior colliculus section was imaged by locating the superior colliculus and then moving 800 µm caudal to where the nucleus was no longer visible. For each section, twenty-three spin-echo images (echo time: 11 ms with a flip angle of 180°, acquisition time: 7.76 ms, voxel size 280×104×400 µm; slice thickness 400 µm; base resolution of 192 mm; field of view 20 mm) were acquired at different repetition times in the following order: 0.15, 3.50, 1.00, 1.90, 0.35, 2.70, 0.25, and 0.50 s. Once animals regained consciousness and were fully ambulatory, they were returned to their home cage.

### 2.6. Image processing and defining regions of interest (ROIs)

Outlines of the ROIs were manually drawn bilaterally onto T1 maps (ImageJ software - Version 1.48p, Wayne Rasband National Institutes of Health, USA) using the Paxinos and Watson rat atlas as a reference. These ROIs consisted of VNC subdivisions: lateral vestibular nucleus (LVe), medial vestibular nucleus - magnocellular (MVeMC), medial vestibular nucleus - parvocellular (MVePC), spinal vestibular nucleus (SpVe), and superior vestibular nucleus (SuVe) and VeCb nuclei: flocculus, nodulus, paraflocculus, and uvula. The rostral VNC included the LVe, MVeMC, MVePC, SuVe, and SpVe (Fig. 2A - B), while the mid rostro-caudal and caudal VNC included MVeMC, MVePC and SpVe (Fig. 2C - F). The paraflocculus of the VeCb was evaluated rostro-caudally. The three other VeCb nuclei, PnO, and Ent nuclei were not divided rostro-caudally.

**Figure 2.**
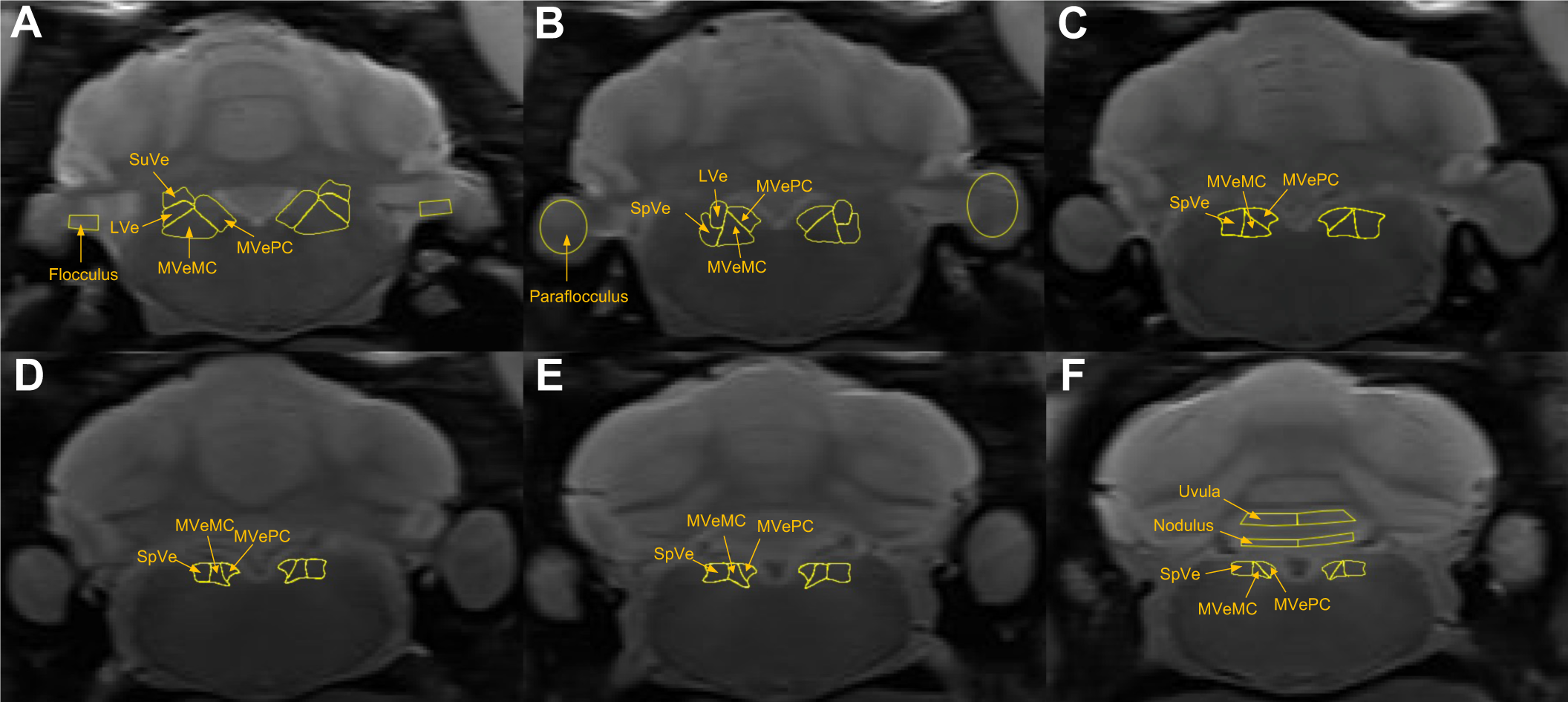
Serial rostro-caudal MRI images through the VNC and VeCb of the rat brain. Average grayscale images were used to outline rostral (A - B), mid rostro-caudal (C – D) and caudal (E – F) nuclei of the VNC and corresponding cerebellar regions. LVe = lateral vestibular nucleus; MVeMC = medial vestibular nucleus, magnocellular division; MVePC = medial vestibular nucleus, parvocellular division; SpVe = spinal vestibular nucleus; SuVe = superior vestibular nucleus.

### 2.7. Statistical analysis

The reciprocal of T1 (1/T1 or *R*1) values from the MRI T1 maps were then used to quantify the extent of Mn^2+^ uptake. T1 images were compared across groups using StatView Version 5.0.1.0 (SAS Institute Inc.). Group differences were analyzed by one-way or two-way analyses of variance (ANOVAs) with Tukey/Kramer post-hoc comparisons. *P* values less than 0.05 were considered significant. F tests were performed to determine whether linear or quadratic functions were the best fit of *R*1 data as a function of rostro-caudal position. Importantly, left and right data for each brain region were averaged in each animal because no significant effect of side on *R*1 values was observed. All data are expressed as mean ± standard error of the mean (SEM).

## 3. Results

### 3.1. Mn^2+^ is taken up and cleared over time from the VNC and VeCb

A major benefit of Mn^2+^ as an MRI contrast agent is slow cellular efflux. This ability of cells to retain Mn^2+^ for hours to days allows a longer time for visualization of encoded neuronal activity. In the current study, baseline MRI *R*1 values were compared with MRI *R*1 values from the VNC, one and 14 days after Mn^2+^ administration (Fig. 3A - I). Comparisons were made in the rostral, mid rostro-caudal, and caudal VNC. In the rostral VNC, *R*1 values were significantly increased one day after Mn^2+^ administration (Fig. 3J). when compared to baseline *R*1 values (baseline: 0.470 ± 0.004; one day: 0.696 ± 0.028; *p*<0.05). By two weeks, *R*1 values (0.494 ± 0.013) were not significantly different from baseline values. The significant increase in *R*1 values after Mn^2+^ administration in the one day group was observed in the mid rostro-caudal (baseline: 0.481 ± 0.009; one day: 0.791 ± 0.051; two weeks: 0.500 ± 0.016, Fig. 3K) (*p*<0.05), and caudal VNC (baseline: 0.465 ± 0.006; one day: 0.729 ± 0.049; two weeks: 0.502 ± 0.018, Fig. 3L) (*p*<0.05). This consistent significant increase in *R*1 (sec^-1^) values in the one-day group, is suggestive of Mn^2+^ uptake throughout the rostro-caudal extent of the VNC.

**Figure 3.**
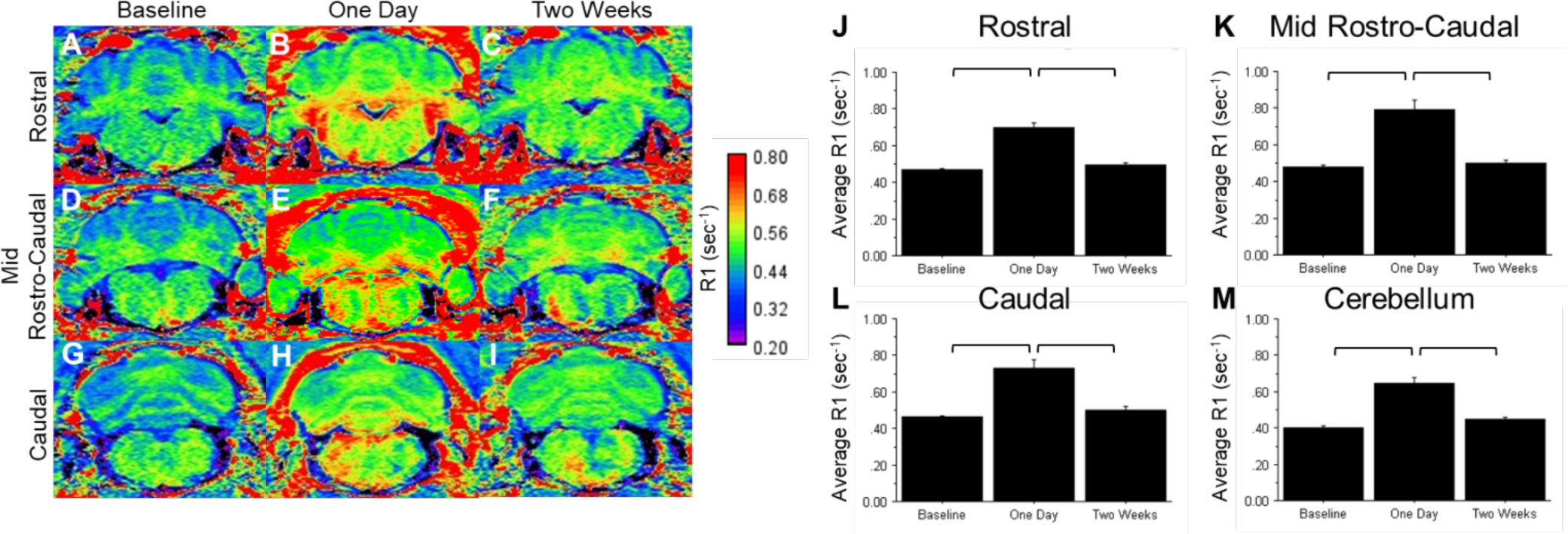
Representative R1 maps from a stimulated rat. In rostral (A – C), mid rostro-caudal (D – F), and caudal (G – I) VNC regions Mn^2+^ uptake was assessed, one day (B, E, H), and two weeks (C, F, I) after administration of MnCl_2_ and compared to baseline (A, D, G). R1 values were compared among the baseline, one day, and two weeks groups in the rostral (J), mid rostro-caudal (K), and caudal (L) VNC, as well as the VeCb (M) for all rats. Colorimetric bar: quantitative R1 scale. Brackets denote significant difference using Tukey’s HSD test for post-hoc analysis (*p*<0.05). Each bar represents the mean ± SEM. For all regions baseline, n=4; one day after Mn^2+^: rostral and mid-rostro-caudal VNC, n=10; caudal VNC, n=8; and cerebellum, n=10; for all regions two weeks after Mn^2+^, n=4.

When *R*1 (sec^-1^) values from the one day group were evaluated two weeks after Mn^2+^ administration, there was a significant decreased in the (rostral: one day: 0.696 ± 0.028; two weeks: 0.494 ± 0.013, mid rostro-caudal: one day: 0.791 ± 0.051; two weeks: 0.500 ± 0.016, caudal: one day: 0.729 ± 0.049; two weeks: 0.502 ± 0.018, Fig. 3J – L, *p*<0.05). However, baseline and two-week *R*1 (sec^-1^) values were not significantly different from one another (rostral: baseline: 0.470 ± 0.004; two weeks: 0.494 ± 0.013, mid rostro-caudal: baseline: 0.481 ± 0.009; two weeks: 0.500 ± 0.016, caudal: baseline: 0.465 ± 0.006; two weeks: 0.502 ± 0.018, Fig. 3J - L). These significant changes in *R*1 (sec^-1^) values after two weeks suggest that Mn^2+^ was cleared from the VNC within two weeks and resulted in reduced *R*1 (sec^-1^) values.

The results were similar for the VeCb. One day after Mn^2+^ administration (Fig. 3M), one day *R*1 values were significantly increased when compared to baseline VeCb *R*1 values (baseline: 0.402 ± 0.009; one day: 0.645 ± 0.032). By two weeks, *R*1 values (0.449 ± 0.012), were not significantly different from baseline values (*p*>0.05). Similar to the VNC, these results suggest that Mn^2+^ is taken up and cleared from the VeCb within two weeks.

### 3.2. Uptake of Mn^2+^ in the VNC is greatest in the mid rostro-caudal regions

For the VNC, baseline *R*1 values across all animals ranged from 0.454 - 0.537 sec^-1^ throughout the 2,000 µm of the VNC that were examined. Baseline *R*1 values across the six sections (0 - 2,000 µm) were best fit to a line and no effect of rostro-caudal position was observed (Fig. 4A, p>0.05). One day after administration, the data from both groups were best fit by quadratic equations, suggesting greater Mn^2+^ uptake in mid rostro-caudal VNC sections (Fig. 4A, Table 1b, sham: *p*=0.050, stimulated: *p*=0.001). After two weeks, the sham group’s *R*1 values were again best fit by a line, with no rostro-caudal effects on *R*1 values. However, the two week stimulated group’s *R*1 values were still better fit by a quadratic than a line (Fig. 4A, *p*=0.039)

**Figure 4.**
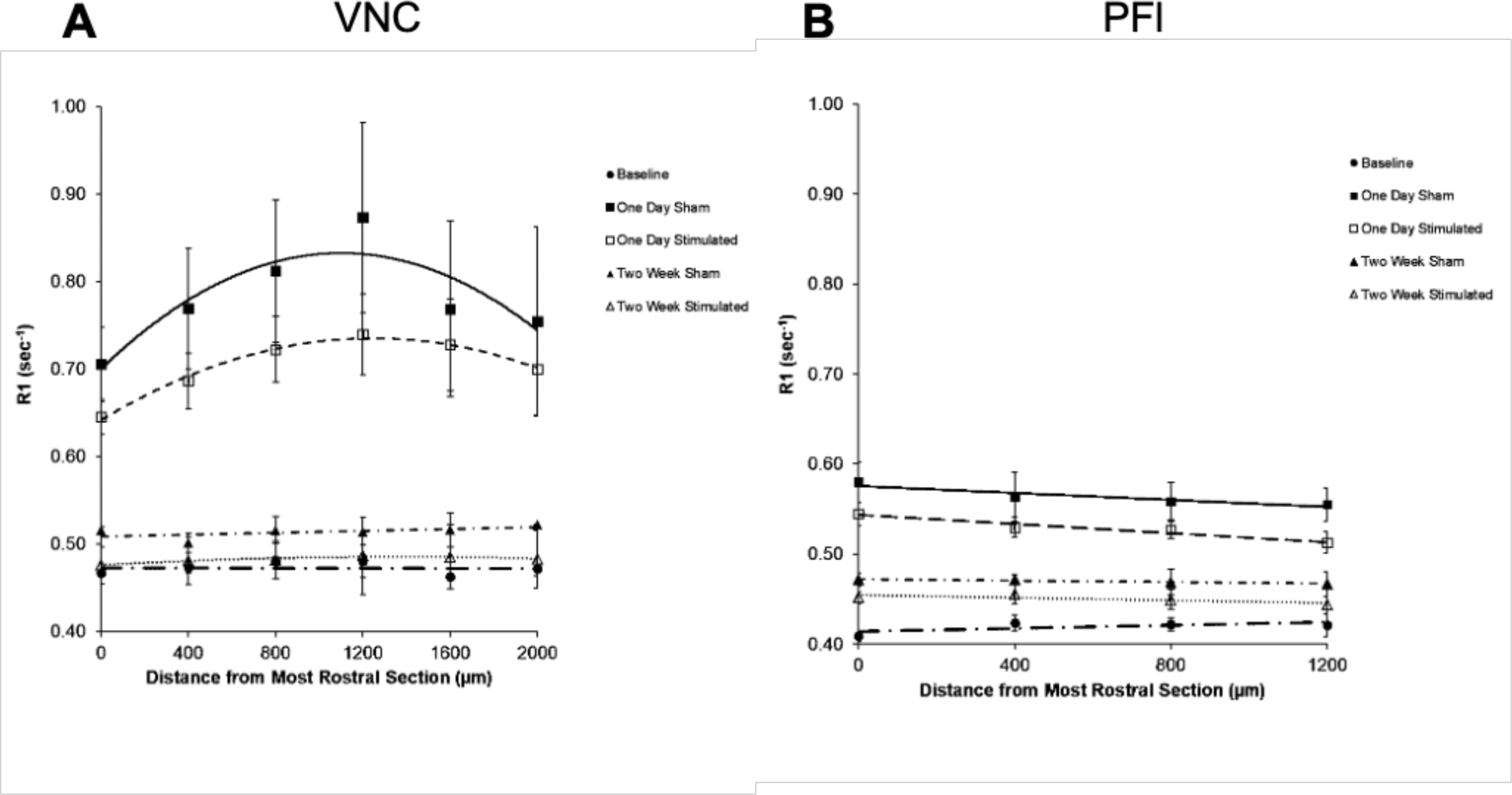
Mn^2+^ uptake across the rostro-caudal extent of the VNC and paraflocculus. Averaged R1 values from six and four serial sections through the VNC and paraflocculus, respectively, are arranged from rostral to caudal. 0 µm corresponds to figure 2A, 400 µm corresponds to figure 2B, etc. Quadratic or linear trend lines are included for each condition depending on which function fit best. Each point represents the mean ± SEM. VNC: Baseline: y = −3.57 × 10^−7^x + 0.48; One Day Sham: y = −1.09 × 10^−7^x^2^ + 2.40 × 10^−4^x + 0.70; One Day Stimulated: y = −6.00 × 10^−8^x^2^ + 1.50 × 10^−4^x + 0.64; Two Week Sham: y = 5.00 × 10^−6^x + 0.51; Two Week Stimulated: y = −5.19 × 10^−9^x^2^ + 1.41 × 10^−5^x + 0.48. Paraflocculus: Baseline: y = 8.75 × 10^−6^x + 0.41; One Day Sham: −1.98 × 10^−5^x + 0.58; One Day Stimulated: −2.55 × 10^−5^x + 0.54; Two Week Sham: y = −4.50 × 10^−6^x + 0.47; Two Week Stimulated: y = −7.75 × 10^−6^x + 0.45. y = R1; x = distance from most rostral section (µm).

**Table 1.** Mean R1 values in each section of the VNC and paraflocculus one day after Mn^2+^ administration. VNC = vestibular nuclear complex; PFl = paraflocculus. Each value represents the mean R1 value ± SEM; n = animals per group

| Distance From Most Rostral Section ($\mu\text{m}$) | VNC<br>Sham: One Day<br>Average R1 ($\text{sec}^{-1}$) | VNC<br>Stimulated: One Day<br>Average R1 ($\text{sec}^{-1}$) | PFI<br>Sham: One Day<br>Average R1 ($\text{sec}^{-1}$) | PFI<br>Stimulated: One Day<br>Average R1 ($\text{sec}^{-1}$) |
| --- | --- | --- | --- | --- |
| **a** 0 (n=5) | 0.705 ± 0.043 | 0.645 ± 0.020 | 0.580 ± 0.022 | 0.545 ± 0.013 |
| **a** 400 (n=4-5) | 0.769 ± 0.069 | 0.686 ± 0.032 | 0.563 ± 0.027 | 0.530 ± 0.011 |
| **b** 800 (n=5) | 0.812 ± 0.081 | 0.722 ± 0.037 | 0.559 ± 0.020 | 0.527 ± 0.009 |
| **b** 1,200 (n=4-5) | 0.873 ± 0.109 | 0.740 ± 0.046 | 0.555 ± 0.018 | 0.512 ± 0.012 |
| **c** 1,600 (n=4) | 0.769 ± 0.100 | 0.728 ± 0.052 | - | - |
| **c** 2,000 (n=3-4) | 0.755 ± 0.108 | 0.700 ± 0.053 | - | - |

a. Rostral sections
b. Mid Rostro-caudal section
c. Caudal sections

### 3.3. Uptake of Mn^2+^ in the paraflocculus is greatest in the rostral regions

For the paraflocculus, baseline *R*1 values across all animals ranged from 0.386 - 0.449 sec^−1^. Baseline *R*1 values across the four sections (0 – 1,200 µm) were best fit to a line and showed no effect of rostro-caudal position, i.e., the slopes were not significantly different from zero (*p*=0.310, Fig. 4B). At one day, the data were also best fit to a line for both groups. The stimulated group had a negative slope that was statistically significant, i.e., *R*1 decreased more caudally (*p*=0.027). The sham group showed a similar negative trend, but the slope only bordered on statistical significance (*p*=0.073). Thus, one day after stimulation, rostral regions took up more Mn^2+^ compared to caudal regions (Fig. 4B, Table 1a-b). After two weeks, the data were again best fit to a line for both groups, with no significant differences in *R*1 values across the rostro-caudal extent of the VeCb.

### 3.4. Jerk stimulation reduces Mn^2+^ uptake in the rostral VNC

Mean *R*1 values were collected from *R*1 maps of each VNC subdivision and compared between the sham and stimulated rats in rostral (Fig. 5A-B), mid rostro-caudal (Fig. 6A-B), and caudal (Fig. 7A-B) sections one day after Mn^2+^ administration. *R*1 values from the rostral VNC showed a significant effect of stimulation in the rostral VNC (Fig. 5C, p=0.026), but no significant effect of stimulation was observed in the mid rostro-caudal or caudal VNC (Figs. 6C and 7C) (**Rostral:** LVe - sham 0.718 ± 0.052; LVe – stimulated, 0.661 ± 0.025; MVeMC – sham, 0.675 ± 0.031; MVeMC – stimulated, 0.626 ± 0.018; MVePC – sham, 0.818 ± 0.086; MVePC – stimulated, 0.733 ± 0.038; SpVe – sham, 0.690 ± 0.042; SpVe – stimulated, 0.624 ± 0.022; SuVe – sham, 0.732 ± 0.051; SuVe – stimulated, 0.668 ± 0.024. **Mid rostro-caudal:** MVeMC, sham - 0.815 ± 0.092; MVeMC – stimulated: 0.727 ± 0.036; MVePC – sham, 0.930 ± 0.131; MVePC – stimulated, 0.793 ± 0.051; SpVe – sham, 0.821 ± 0.082; SpVe – stimulated, 0.715 ± 0.032. **Caudal:** MVeMC – sham, 0.729 ± 0.093; MVeMC – stimulated, 0.707 ± 0.057; MVePC – sham: 0.787 ± 0.111; MVePC – stimulated: 0.757 ± 0.072; SpVe – sham, 0.726 ± 0.075; SpVe - stimulated, 0.707 ± 0.044).

**Figure 5.**
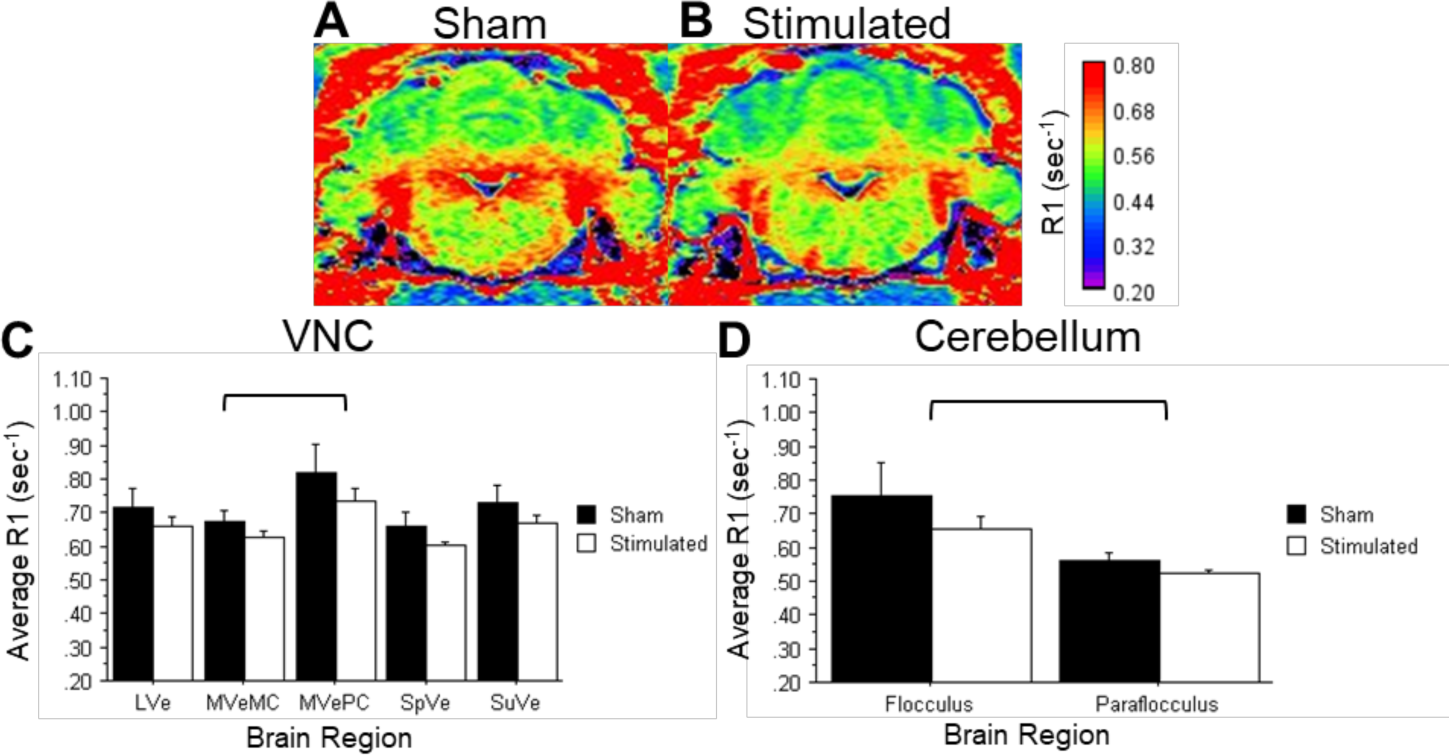
Mn^2+^ uptake in rostral VNC sub-nuclei and VeCb one day after Mn^2+^ administration in sham and stimulated animals. R1 maps show Mn^2+^ uptake in the rostral VNC and corresponding cerebellar regions in a sham (A) and stimulated (B) rat. Corresponding R1 values were compared between sham (n=5) and stimulated (n=5) rats in rostral VNC (C) and cerebellar nuclei (D). LVe = lateral vestibular nucleus; MVeMC = medial vestibular nucleus, magnocellular division; MVePC = medial vestibular nucleus, parvocellular division; SpVe = spinal vestibular nucleus; SuVe = superior vestibular nucleus. Colorimetric bar: quantitative R1 map scale. Two-way ANOVA analysis of panels C and D showed a significant impact of brain region (p<0.05) and stimulation (p<0.05). Brackets denote significant differences between brain regions determined by Tukey HSD post-hoc analysis. Each bar represents the mean R1 value ± SEM.

**Figure 6.**
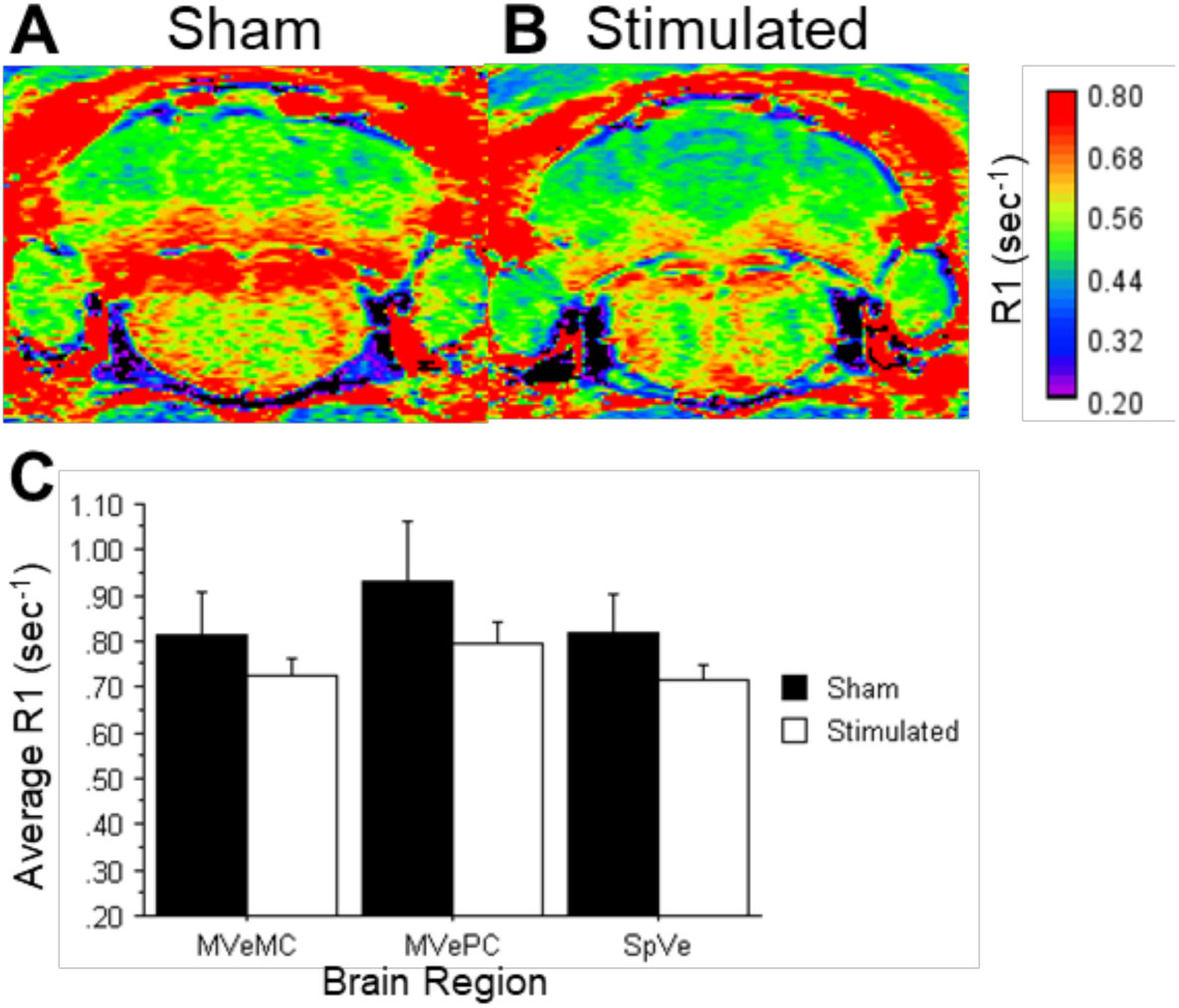
Mn^2+^ uptake in mid rostral-caudal VNC sub-nuclei one day after Mn^2+^ administration. R1 maps through the mid rostro-caudal VNC are shown for a sham (A) and stimulated (B) rat. The corresponding R1 values were compared between sham (n=5) and stimulated (n=5) rats in mid rostro-caudal VNC subnuclei (C). MVeMC = medial vestibular nucleus, magnocellular division; MVePC = medial vestibular nucleus, parvocellular division; SpVe = spinal vestibular nucleus. Colorimetric bar: quantitative R1 scale. Each bar represents the mean ± SEM.

**Figure 7.**
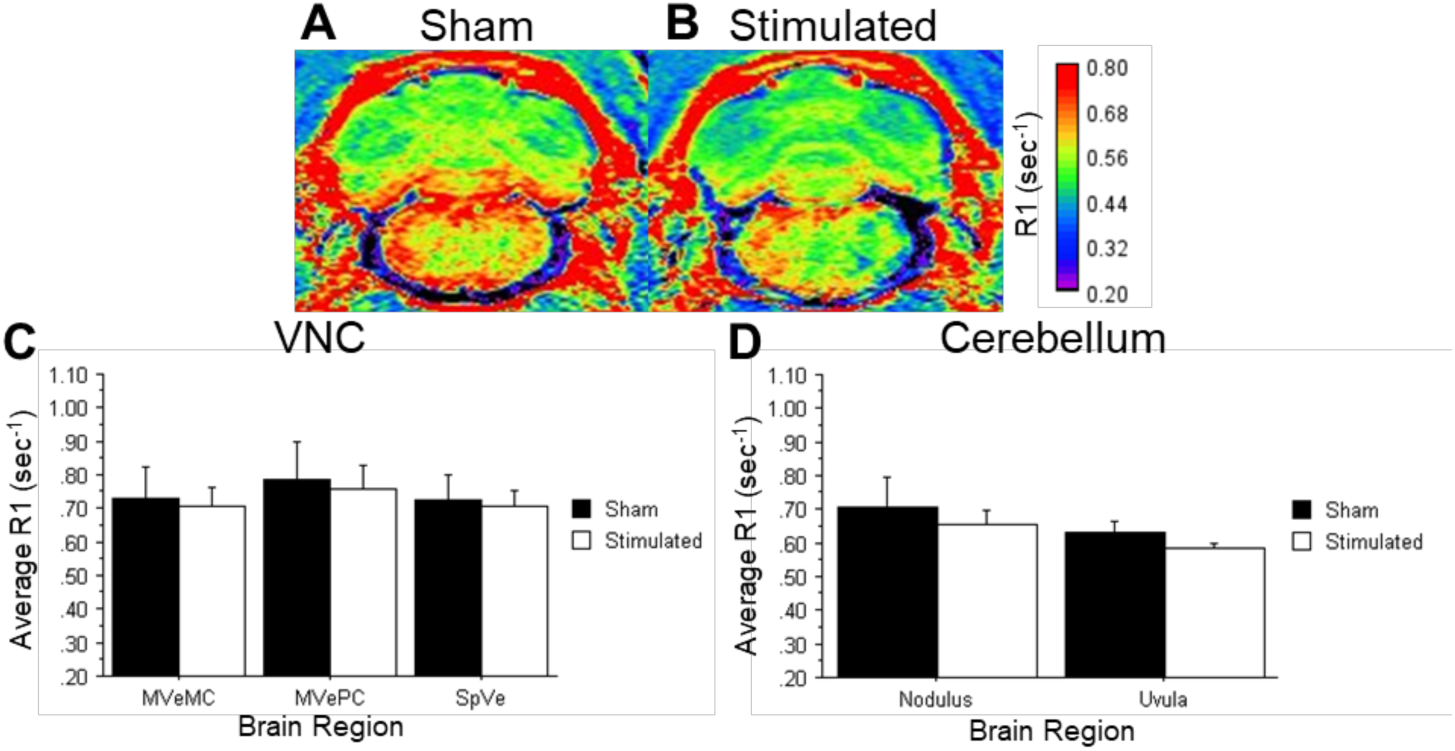
Mn^2+^ uptake in caudal VNC sub-nuclei and VeCb one day after Mn^2+^ administration. R1 maps through the caudal VNC are shown for a sham (A) and stimulated (B) rat. The corresponding R1 values were compared between sham (n=4) and stimulated (n=4) rats in the caudal VNC subnuclei (C) and cerebellar nuclei (D). MVeMC = medial vestibular nucleus, magnocellular division; MVePC = medial vestibular nucleus, parvocellular division; SpVe = spinal vestibular nucleus. Colorimetric bar: quantitative R1 scale. Each bar represents the mean ± SEM.

### 3.5. Mn^2+^ is differentially taken up within rostral VNC subdivisions

A significant effect of brain region (VNC subdivisions) was observed in the rostral (Table 1, 0 - 400 µm) VNC (Fig. 5C; *p*=0.047). There was significantly less Mn^2+^ uptake in the MVeMC (0.650 ± 0.019) compared to the MVePC (0.776 ± 0.047; *p*<0.05) in the rostral sections. No significant effect of brain region was observed in the mid rostro-caudal and caudal VNC sections (Fig. 6C and 7C – D).

### 3.6. Stimulation does not affect Mn^2+^ uptake in the VeCb

No effect of stimulation was observed in rostral and caudal VeCb regions (**Rostral VeCb**: flocculus – sham, 0.754 ± 0.097; flocculus – stimulated, 0.654 ± 0.036; paraflocculus – sham, 0.563 ± 0.021; paraflocculus – stimulated, 0.524 ± 0.009. **Caudal VeCb:** nodulus – sham, 0.708 ± 0.086; nodulus – stimulated, 0.655 ± 0.043; uvula – sham, 0.629 ± 0.037; uvula – stimulated, 0.585 ± 0.015).

### 3.7. Mn^2+^ is differentially taken up in the VeCb

In rostral VeCb regions (flocculus and paraflocculus, Fig. 2 and 5A-B), there was a significant effect of brain region on *R*1 values (Fig. 5D, *p*=0.008). The flocculus had significantly greater *R*1 values than the paraflocculus (*p*<0.05, flocculus: 0.704 ± 0.052, paraflocculus: 0.543 ± 0.012). In the caudal VeCb there was no effect of brain region when comparing the nodulus and uvula (Fig. 7, nodulus: 0.681 ± 0.046, uvula: 0.607 ± 0.020).

### 3.8. Stimulation does not affect Mn^2+^ uptake in non-vestibular related brain regions

As control brain regions, we looked at *R*1 values using the *R*1 maps for two brain regions that are not traditionally considered part of the vestibular pathway (Fig. 8A-B). In both the entorhinal cortex (Ent) and the pontine reticular nucleus, oral part (PnO), there were no differences between the sham and stimulated groups (Fig. 8C, *p*=0.753).

**Figure 8.**
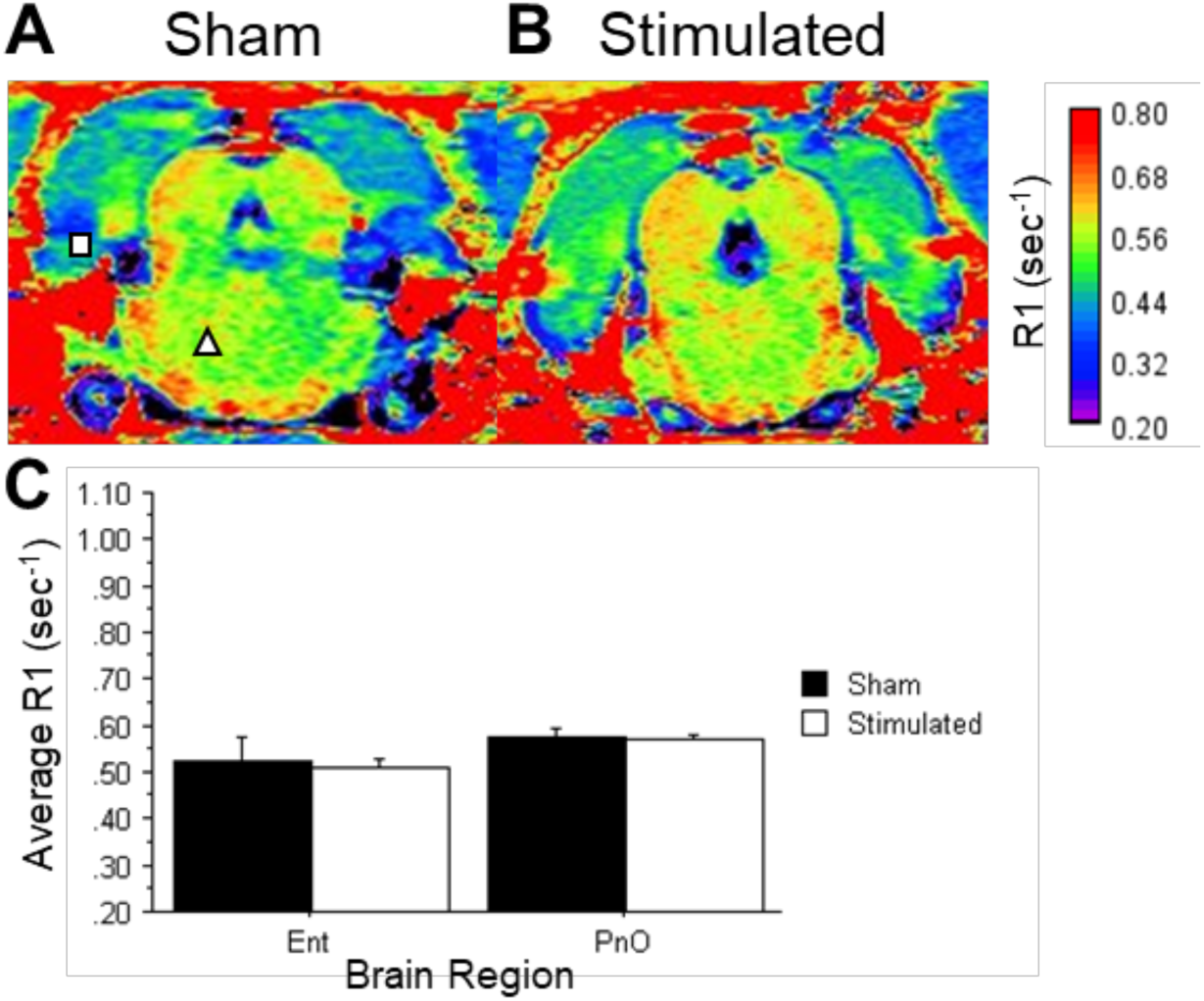
Mn^2+^ uptake in non-vestibular regions one day after Mn^2+^ administration. R1 (sec^−1^) maps depict analyzed regions from the entorhinal cortex - square and pontine reticular nucleus - triangle. The corresponding R1 values were compared between sham (n=4) and stimulated rats (n=5). Ent = entorhinal cortex; PnO = pontine reticular nucleus, oral part. Colorimetric bar: quantitative R1 scale. Each bar represents the mean ± SEM.

## 4. Discussion

### 4.1. MEMRI can be used to assess neuronal activity in the VNC and VeCb

To our knowledge, this is the first study to successfully show that each of the VNC and VeCb subdivisions can take up Mn^2+^ following administration of MnCl_2_ (Fig. 3). Although a previous study has shown Mn^2+^ uptake in the medial vestibular nucleus in mice administered MnCl_2_ (Rodriguez et al., 2016), they did not report the status of Mn^2+^ uptake for each of the VNC subdivisions. Since our data show differential *R*1 values among VNC and VeCb subdivisions following MnCl_2_ administration, this implies that these brain regions take up Mn^2+^ in an activity-dependent manner. Uptake of Mn^2+^ was evident throughout rostro-caudal regions of interest for both the VNC and VeCb (Fig. 3). Also, our data show that Mn^2+^ clears from these active regions within two weeks after administration. As a result, additional MEMRI assessments may be conducted in the same animal over time, providing the benefit of repetitive *in vivo* measurement of central vestibular activity.

### 4.2. Otolith stimulation and localization of central neuronal activity

Due to the orientation of the animal’s head, we believe the otolith organs are the primary contributors to VNC and VeCb activity patterns in both sham and stimulated rats. The jerk stimulus delivered in the current study is a form of linear acceleration, which when delivered in the naso-occipital plane (dorso-ventral), predominantly activates the saccule. Regardless of stimulation, tonic information is detected by hair cells of the otolith organs with respect to gravity (Feher, 2012). Since the force of gravity is present, we expect spontaneous activity of the otolith organs to contribute to activity patterns observed in the sham rats. Although the linear acceleration in the naso-occipital plane and head orientation of the animals implicate the saccule as the primary end-organ being stimulated in our study, the asymmetric and curved nature of otolith maculae suggests that the utricle may also contribute to the response. The striola of these saccular and utricular otolith organs are innervated by rapidly adapting irregular fibers that terminate in the VNC (Desmadryl & Dechesne, 1992; Eatock et al., 2008; Goldberg et al., 1990). Furthermore, since rotational head movements were minimal, contributions from the semicircular canals were not expected. Therefore, this model of bidirectional jerk stimulation that preferentially activates irregular fibers provides a reliable method for understanding relationships between peripheral otolith stimulation and central activation of secondary neurons in the VNC and cerebellar circuitry.

### 4.3. Rostro-caudal Mn^2+^ uptake in the VNC corresponds with vestibulo-autonomic pathways

The vestibular system can have direct effects on autonomic centers modulating affect, heart rate, and blood pressure. These functions are segregated within the VNC, with affect, emotion, and modulation of intestinal viscera in rostral subdivisions (MVe and SuVe) (Balaban, 1996; Balaban & Yates, 2004; Kerman et al., 2006) and blood pressure and heart rate in more caudal subdivisions (MVe and SpVe; Holstein, 2012; Balaban and Yates 2004). Otolith projections generally target more caudal VNC regions compared to rostral VNC regions (Dickman and Angelaki, 2002; Newlands and Perachio, 2003). These mid rostro-caudal to caudal VNC neurons receiving otolith projections send direct and indirect projections to autonomic brainstem nuclei, including the nucleus tractus solitarius (NTS) and the rostral ventrolateral medulla (RVLM; Balaban and Beryozkin, 1994; Porter and Balaban, 1997; Yates et al., 2013), regions which affect blood pressure and heart rate. Semi-circular canals predominantly target rostral VNC regions. These preferential projections and functions support the more active regions (greater Mn^2+^ uptake) we observed in the more caudal regions of the VNC (Fig. 4), regardless of the stimulation status of the animal. The mid rostro-caudal VNC regions delineated in the present study may partially encompass regions considered caudal in previous studies. The designations of rostral or caudal have traditionally been applied to VNC regions (Holstein, 2012). The regions designated as mid rostro-caudal in the current study primarily contain VNC subdivisions found more caudally (MVe and SpVe), with no LVe region present (Fig. 2).

### 4.4. Repetitive mild linear acceleration reduces activity in the rostral VNC

Although the pattern of activation in VNC subnuclei was similar in sham and stimulated animals, the sham animals showed greater Mn^2+^ uptake when compared to stimulated animals (Fig. 5C). While surprising, there are several potential explanations for these results. A recent study has shown that repeated galvanic vestibular stimulation (GVS) of the vestibular nerve resulted in reduced glutamate receptors, AMPA and NMDA receptors (Kim et al., 2020). This reduction in glutamate receptors may be a compensatory response of the cells to over-excitation and excessive glutamate release. As a result, we may have observed less neuronal activity in the stimulated animals due to a potential reduction in excitatory transmission.

Previous studies have shown that macular primary afferents are spontaneously active in mature animals (Fernández et al. 1972; Goldberg and Fernández 1971; Jones and Jones 2000). The modest stimulation intensity used in the current study (3,000 jerks at ∼ 600 g/s) may reduce Mn^2+^ uptake by producing brief periods of synchronous activity that do not exceed the levels of spontaneous firing that occur when irregular fibers are at rest (Goldberg et al., 1984; Goldberg et al., 1987; Goldberg, 2000). Another possibility for less Mn^2+^ uptake in response to jerk stimulation may be that the stimulus in the present study may inhibit the tonic activity of the regular afferents, reducing the Mn^2+^ uptake in second order vestibular neurons (Goldberg and Fernández, 1971). Our use of a bidirectional jerk stimulus may contribute to greater Mn^2+^ uptake in the sham group. The bidirectional sensitivity of the hair cell bundles that are oriented in opposite directions along the line of polarity reversal (LPR) may account for the reduced activity in apposing irregular fibers (López-Schier et al., 2004; Flock, 1964; Lewis et al., 1985; Fritzsch and López-Schier, 2014; Fritzsch et al., 2002). Stimulation in a particular direction will simultaneously generate an excitatory and inhibitory signal in the otolith maculae which will be transduced to the VNC as well as the uvula and nodulus. Whether stimulation is generating a greater excitatory or inhibitory signal is unclear. However, given the bidirectional sensitivity of these hair cells along the LPR, our data suggests that the mild, repetitive, bidirectional jerk stimuli are largely inhibiting these hair cells which may explain the reduced activity we observed in the VNC and VeCb.

Cross-striolar inhibition is another mechanism that may account for the observed activity patterns in which VNC neurons on one side are disynaptically inhibited by contralateral afferents (Uchino et al., 1997). This mechanism increases the sensitivity of vestibulo-spinal neurons to bilateral inputs (Uchino and Kushiro, 2011) as there is convergence of facilitation from the ipsilateral side and disinhibition from the contralateral side. Since our stimulation is bilateral, cross-striolar inhibition may explain why stimulated rats show less activity in the VNC than sham rats since both sides of the VNC are being di-synaptically inhibited by the contralateral afferents. The reduced activity we observed may be due to the repeated jerk stimulation (Fig. 1D).

Our data showing that the greatest Mn^2+^ uptake in the MVePC were unexpected, given that the SpVe and LVe are enriched for projections from otolith organs (Büttner-Ennever, 1992; Barmack, 2003; Highstein and Holstein, 2006; Goldberg et al., 2012). However, our results (Fig. 5C) were similar to a previous study, in which sinusoidal galvanic vestibular stimulation (sGVS) of the 8^th^ nerve, resulted in the greatest density of immunolabeling for a marker of neuronal activity, c-Fos, in the MVePC (Holstein et al., 2012). In this same study, c-Fos immunolabeling was largely absent in the MVeMC.

### 4.5. Differential activity patterns are observed across vestibular subnuclei

Unlike the flocculus and paraflocculus, both the nodulus and uvula receive direct afferent input from the vestibular nerve; axons originating from the saccule directly target the uvula whereas canal input targets the nodulus (Maklad and Fritzsch, 2003; Newlands et al. 2003). Our results demonstrating greater Mn^2+^ uptake in the flocculus compared to the paraflocculus (Fig. 5D) and no significant difference in Mn^2+^ uptake between the nodulus and uvula were unexpected based upon known projections and the stimulation that we delivered. Again, while the modest 3,000 jerks delivered at ∼600 g/s in the present study was capable of changing Mn^2+^ uptake in the rostral VNC, the question of whether intensity dependent changes in uptake would be observed with increasing stimulus intensity remains unknown. Also, whether the effect of changes in the intensity or number of jerks delivered would be equal across VNC subdivisions would be fruitful for future studies. Using MEMRI as an *in vivo* marker of neuronal activity, future studies could also be developed to examine differences in activity patterns among VNC and VeCb regions comparing linear acceleration to angular acceleration stimulation of irregular fibers.

### 4.6. Linear acceleration has no effect on neuronal activity in non-vestibular related brain regions

As control brain regions, we assessed Mn^2+^ uptake in the PnO and the Ent since we do not expect to see a difference between sham and stimulated groups. The oral part of the pontine reticular nucleus is involved with forming the paradoxical stage of sleep (Garzón et al., 1998; Rechtschaffen and Siegel 2000; Xi et al., 1999) and has a significant role in sleep disorders when lesioned (J. Arpa, 1995). Since it has not been shown to be involved with the vestibular system, we looked at the Mn^2+^ uptake and saw no difference between the sham and stimulated group. Moreover, we also looked at the Ent, which is involved with learning and memory. It showed a similar pattern as the PnO, showing no significant difference between treatment groups. This supports our results that the difference in Mn^2+^ uptake between groups is not due to potential confounding variables, (i.e., differences in Mn^2+^ availability in the blood), but differences in neuronal activity in the VNC and VeCb in response to otolith irregular fiber stimulation.

### 4.7. Future implications of MEMRI application in vestibular pathways

Due to the activity-dependent changes in Mn^2+^ uptake we observed in the VNC and VeCb, our data suggest that MEMRI may be applied in future animal studies to investigate changes in central neuronal activity and plasticity following vestibular injury, as previously seen with blast (Rodriguez et al., 2016). Our data showing clearance of Mn^2+^ from the brain regions by two weeks after MnCl_2_ administration also support the possibility of longitudinal tracking of changes in neuronal activity across vestibular subdivisions in the same animal. Additionally, our pre-clinical studies using MEMRI pave the way for future clinical studies evaluating diagnostic tools and treatments for central vestibular disorders.

## Acknowledgments

We thank Aaron K. Apawu, Srinivasu Kallakuri, and Tasnia Chowdhury, for contributions to the data analysis and to Sharowynn Wilson for contributions to data organization and pre-manuscript review. We also thank the John D. Dingell VAMC and Wayne State University School of Medicine for their ongoing support of this research. Funding of these experiments was made possible by the Department of Veterans Affairs (Grant 1I01RX001095 U.S to A.G.H); Department of Veterans Affairs (Grant 1I01RX001986 to A.G.H). The views expressed in this paper do not necessarily reflect the official policies of the Department of Health and Human Services, nor does mention of trade names, commercial practices, or organizations imply endorsement by the U.S. Government.

